# Exogenous alginate protects *Staphylococcus aureus* from killing by *Pseudomonas aeruginosa*

**DOI:** 10.1101/748459

**Authors:** Courtney E. Price, Dustin G. Brown, Dominique H. Limoli, Vanessa V. Phelan, George A. O’Toole

**Affiliations:** Department of Microbiology and Immunology, Geisel School of Medicine at Dartmouth; Department of Pharmaceutical Sciences, Skaggs School of Pharmacy and Pharmaceutical Sciences, University of Colorado, Aurora, CO; Department of Microbiology and Immunology, University of Iowa, Iowa City, IA

**Keywords:** alginate, cystic fibrosis, *Pseudomonas aeruginosa*, *Staphylococcus aureus*, polymicrobial, mucoid, Nanostring, PQS, HQNO, pyoverdine, pyochelin, siderophores

## Abstract

Cystic fibrosis (CF) patients chronically infected with both *Pseudomonas aeruginosa* and *Staphylococcus aureus* have worse health outcomes than patients who are mono-infected with either *P. aeruginosa* or *S. aureus*. We showed previously that mucoid strains of *P. aeruginosa* can co-exist with *S. aureus in vitro* due to transcriptional downregulation of several toxic exoproducts typically produced by *P. aeruginosa*, including siderophores, rhamnolipids, and HQNO (2-heptyl-4-hydroxyquinoline N-oxide). Here we demonstrate that exogenous alginate protects *S. aureus* from *P. aeruginosa* in both planktonic and biofilm co-culture models under a variety of nutritional conditions. *S. aureus* protection in the presence of exogenous alginate is due to transcriptional downregulation of *pvdA*, a gene required for the production of the iron scavenging siderophore pyoverdine, as well as down-regulation of the PQS (Pseudomonas quinolone signal; 2-heptyl-3,4-dihydroxyquinoline) quorum sensing system. The impact of exogenous alginate is independent of endogenous alginate production. We further demonstrate that co-culture of mucoid *P. aeruginosa* with non-mucoid *P. aeruginosa* can mitigate the killing of *S. aureus* by the non-mucoid strain of *P. aeruginosa*, indicating that the mechanism we describe here may function *in vivo* in the context of mixed infections. Finally, we investigated a panel of mucoid clinical isolates that retain the ability kill *S. aureus* at late time points, and show that each strain has a unique expression profile, indicating that mucoid isolates can overcome the effects of mucoidy in a strain-specific manner.

**IMPORTANCE:** CF patients are chronically infected by polymicrobial communities of microorganisms. The two dominant bacterial pathogens that infect CF patient lungs are *P. aeruginosa* and *S. aureus*, with ∼30% of patients co-infected by both species. Patients infected with both *P. aeruginosa* and *S. aureus* have worse outcomes than mono-infected patients, and both species persist within the same physical space in the lungs of CF patients. A variety of host and environmental factors have been demonstrated to promote *P. aeruginosa*-*S. aureus* co-existence, despite evidence that *P. aeruginosa* kills *S. aureus* when these organisms are co-cultured *in vitro*. Thus, a better understanding of *P. aeruginosa-S. aureus* interactions, particularly mechanisms by which these microorganisms are able to co-exist in proximal physical space, will lead to better informed treatments for chronic polymicrobial infections.

## INTRODUCTION

Cystic fibrosis (CF) is an autosomal recessive disorder with significant morbidity and mortality affecting >70,000 people worldwide (1, 2). Decreased function of the cystic fibrosis transmembrane regulator (CFTR), an epithelial cell chloride ion transporter, causes increased viscosity of airway surface liquid and decreased mucociliary clearance of microorganisms in the CF airway (3, 4). Over time these microorganisms form complex, polymicrobial communities which are highly tolerant to antibiotic treatment; the airway damage due to these chronic infections eventually leads to respiratory failure (5–15). The development of novel therapeutics have markedly improved patient survival (2, 16–19), but despite these recent advances, the progression of CF pulmonary disease is one of chronic infection and inflammation punctuated by periods of clinical exacerbation which cause irreversible damage to lung tissue (2, 8, 20–22).

*Pseudomonas aeruginosa* and *Staphylococcus aureus* are the two opportunistic pathogens most commonly isolated from CF patients’ lungs (2), and co-infection with both microbes is common (23, 24). Co-infection with *P. aeruginosa* and *S. aureus* alters antibiotic tolerance (25–28) and enhances virulence (29) in chronic infection. Furthermore, CF patients who are co-infected with both *P. aeruginosa* and *S. aureus* have poorer clinical outcomes than those who are mono-infected (23).

Studies of *P. aeruginosa*-*S. aureus* interactions have demonstrated that *P. aeruginosa* kills *S. aureus in vitro*, and this interaction is thought to contribute to the dominance of *P. aeruginosa* over *S. aureus* as CF patients age (30, 31). Considerable progress has been made in elucidating the mechanism of *S. aureus* killing mediated by *P. aeruginosa* (32–35). *P. aeruginosa* secretes quorum sensing regulated antimicrobial exoproducts during acute infection, including 2-heptyl-4-hydroxyquinoline N-oxide (HQNO), siderophores, rhamnolipids, and phenazines. HQNO and the two *P. aeruginosa* siderophores, pyoverdine and pyochelin, have been shown to drive *S. aureus* towards a fermentative lifestyle (33), while rhamnolipids disrupt cell membrane integrity (33, 36). Phenazines inhibit *S. aureus* metabolism, as well as play a role in iron acquisition and biofilm development (37–39).

Chronic CF infection is marked by the emergence of mucoid *P. aeruginosa* isolates, which overproduce the exopolysaccharide alginate (40, 41). *P. aeruginosa* virulence towards *S. aureus* is decreased due to the overproduction of alginate, which causes transcriptional downregulation of HQNO, siderophores, and rhamnolipids (32, 33, 42). The mechanism of alginate overproduction is typically due to mutation in the *mucA* gene (43–45), a negative regulator of σ^22^ (AlgT/U) (43, 44, 46). De-repression of σ^22^ promotes transcription of many genes, but disruption of *algD*, the first gene in the alginate biosynthetic operon, restores wild-type levels of antimicrobial exoproducts, demonstrating that alginate production is sufficient for antimicrobial exoproduct downregulation independent of other transcriptional effects of AlgT/U (32).

In this study we asked how alginate influences the production of anti-*Staphylococcal* antimicrobials. We show that exogenous alginate is sufficient to protect *S. aureus* from *P. aeruginosa* in co-culture, likely via the transcriptional downregulation of several genes whose products are required for *P. aeruginosa* to kill *S. aureus*. We also show that a three-way co-culture between mucoid and non-mucoid *P. aeruginosa* and *S. aureus* results in attenuated killing of *S. aureus*, raising the possibility that the mechanism we identified may function *in vivo* in patients with such mixed infections. Furthermore, the analysis of mucoid isolates of *P. aeruginosa* indicates that some isolates are able to kill S*. aureus* after prolonged co-culture and that the mechanism of killing is due to diverse transcriptional changes, indicating a dynamic and evolving competition between these two microbes.

## RESULTS

### Exogenous alginate protects *S. aureus* in co-culture with *P. aeruginosa*

To determine whether the presence of exogenous alginate can promote co-existence between *P. aeruginosa* and *S. aureus*, we investigated a co-culture with 1% seaweed-derived alginate (Fig. 1A-B; Fig. S1). Cultures were plated onto *Pseudomonas* Isolation Agar (PIA) and Mannitol Salt Agar (MSA) to quantify colony forming units (CFUs) of *P. aeruginosa* and *S. aureus*, respectively, over a 10 hour co-culture. Because there are minor structural differences between seaweed-derived alginate and *P. aeruginosa*-produced alginate (46), with seaweed derived alginate lacking acetylation, we also isolated alginate from the mucoid *P. aeruginosa* PAO1 *mucA22* strain (see Materials and Methods) and added this *P. aeruginosa* PAO1-derived alginate to *P. aeruginosa-S. aureus* planktonic cocultures (Fig. 1 A-B; Fig S1). Seaweed-derived and *P. aeruginosa*-derived alginate protected *S. aureus* equally well from killing by *P. aeruginosa* (Fig. 1A; Fig. S1A). 1% seaweed-derived alginate did not affect *P. aeruginosa* growth, while *P. aeruginosa*-derived alginate reduced the growth of *P. aeruginosa* by a small but statistically significant amount (Fig. 1B; Fig. S1B). We performed several additional experiments which supported our conclusion that the addition of exogenous alginate protects *S. aureus* in planktonic coculture with *P. aeruginosa* by a mechanism other than reduced *P. aeruginosa* growth, and expanded our initial results to demonstrate that a range of exogenous alginate (0.25%-2%) concentrations protect *S. aureus* from *P. aeruginosa* in planktonic co-culture (Fig. S2-3).

**Figure 1.**
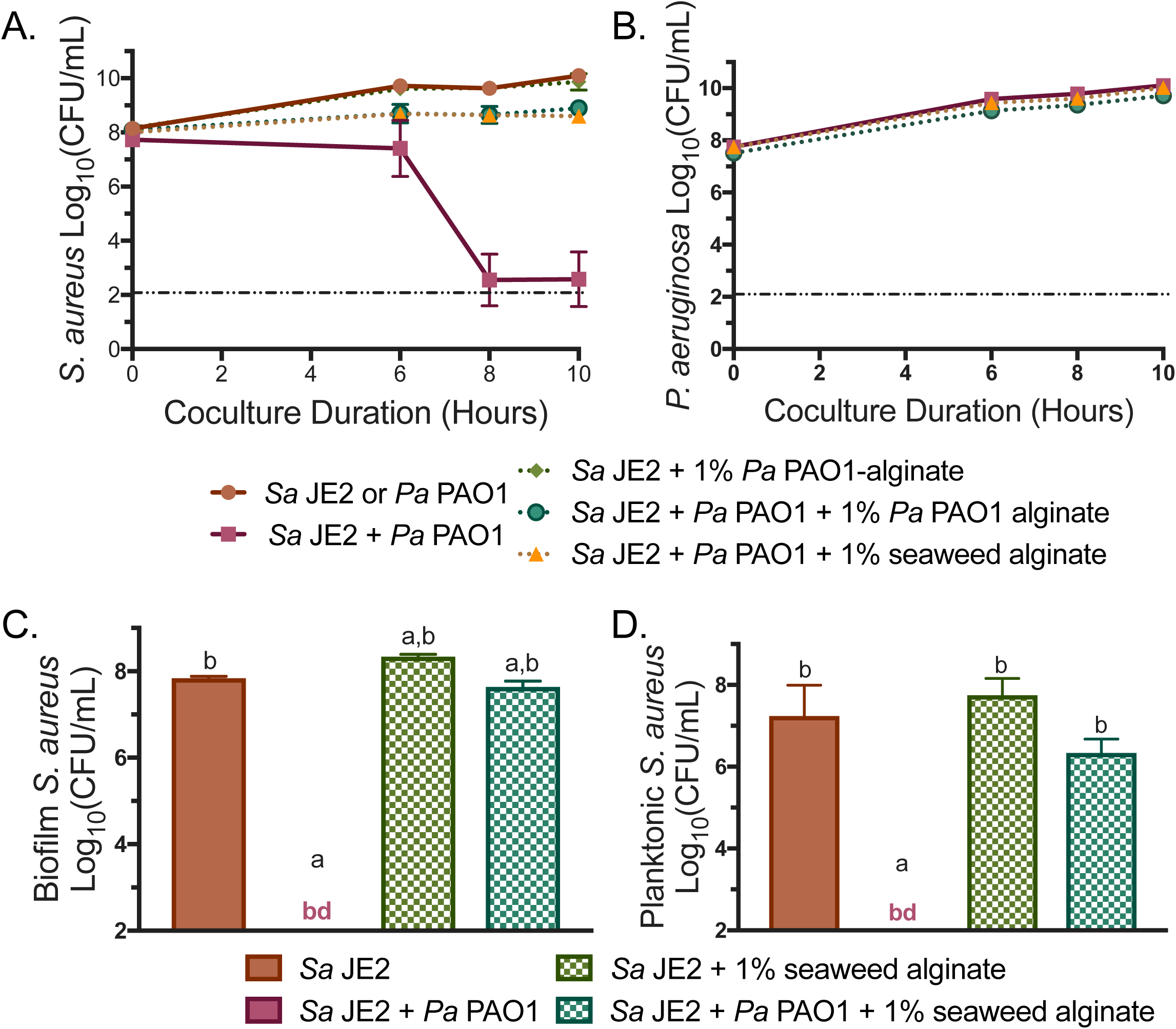
Exogenous alginate protects *S. aureus* JE2 from *P. aeruginosa* PAO1 in co-culture. **A)** *S. aureus* JE2 and **B)** *P. aeruginosa* PAO1 growth curves over 10 hours in liquid co-culture in TSB +/− 1% seaweed-derived or 1% *P. aeruginosa* PAO1-derived alginate. CFU/ml were enumerated at indicated time points and log_10_ transformed. Dotted lines indicate the presence of alginate in the culture. The dashed horizontal line at 2 log_10_ CFU/ml indicates the limit of detection. **C)** Biofilm and **D)** Planktonic *S. aureus* JE2 growth after 16 hours of static co-culture in MEM L-gln L-arg +/− 1% seaweed-derived alginate. CFU/ml were enumerated and log_10_ transformed. Dashed boxes indicate the presence of alginate in the culture. Significance was determined by one-way ANOVA with Dunnett’s post-test. a, p<0.05 with *S. aureus* JE2 as the reference. b, p<0.0001 with *P. aeruginosa* PAO1 + *S. aureus* JE2 as the reference.

Bacterial interactions can change dramatically under different conditions, so we used a coculture model previously described by our lab to determine whether exogenous alginate also protects *P. aeruginosa* from *S. aureus* in a biofilm-based coculture model (25, 33). After an initial 1-hour attachment phase followed by washing to remove planktonic cells, co-cultures of *P. aeruginosa* and *S. aureus* were incubated statically for 16 hours, and then CFUs were enumerated separately for the planktonic and biofilm fractions (see Materials and Methods). Biofilm coculture experiments were performed with *P. aeruginosa* PAO1 (Fig. 1 C-D; Fig. S3 E-F) and *P. aeruginosa* PA14 (Fig. S3, G-J). *S. aureus* was killed by *P. aeruginosa* PAO1 in this biofilm model and protected from *P. aeruginosa* PAO1 with 1% exogenous alginate in both the biofilm (Fig. 1 C) and planktonic (Fig. 1 D) fraction. *P. aeruginosa* PAO1 growth was not affected by exogenous alginate in either fraction (Fig. S3, E-F). Results were similar for *P. aeruginosa* PA14, except that exogenous alginate also modestly boosted *P. aeruginosa* PA14 growth (Fig. S3, I-J).

Experiments with similar results were also performed planktonically with the same minimal media used for biofilm co-culture experiments (Fig. S4). Taken together, these data indicate that exogenous alginate can modulate the killing of *S. aureus* by *P. aeruginosa*, and that exogenous alginate alone is sufficient to protect *S. aureus* from *P. aeruginosa* under a variety of growth conditions.

### Tri-culture of *S. aureus* with wild-type and mucoid *P. aeruginosa* delays *S. aureus* killing

Because exogenous alginate can modify interactions between *P. aeruginosa* and *S. aureus*, and CF patients who are co-infected with *P. aeruginosa* and *S. aureus* are also likely infected with mucoid and non-mucoid *P. aeruginosa* isolates, we hypothesized that mucoid *P. aeruginosa* might affect interactions between wild-type *P. aeruginosa* and *S. aureus*. We therefore designed a tri-culture experiment where *S. aureus* was either monocultured, co-cultured with nonmucoid *P. aeruginosa* PAO1 or mucoid *P. aeruginosa* PAO1 *mucA22*, or cultured with both nonmucoid and mucoid *P. aeruginosa* (Fig. 2; Fig. S5). Each strain of *P. aeruginosa* was inoculated such that experiments with both mucoid and nonmucoid *P. aeruginosa* strains contained 2X the initial concentration of *P. aeruginosa*, to ensure that the total concentration of wild-type *P. aeruginosa* remained consistent across all conditions. *S. aureus* survival increased significantly in both the biofilm and planktonic fractions after 12 hours (Fig. 2A-B) but not after 16 hours (Fig. 2 C-D) of tri-culture with both mucoid and non-mucoid *P. aeruginosa*. These results indicate that mucoid strains can impact interactions between nonmucoid *P. aeruginosa* and *S. aureus* by delaying *S. aureus* killing.

**Figure 2.**
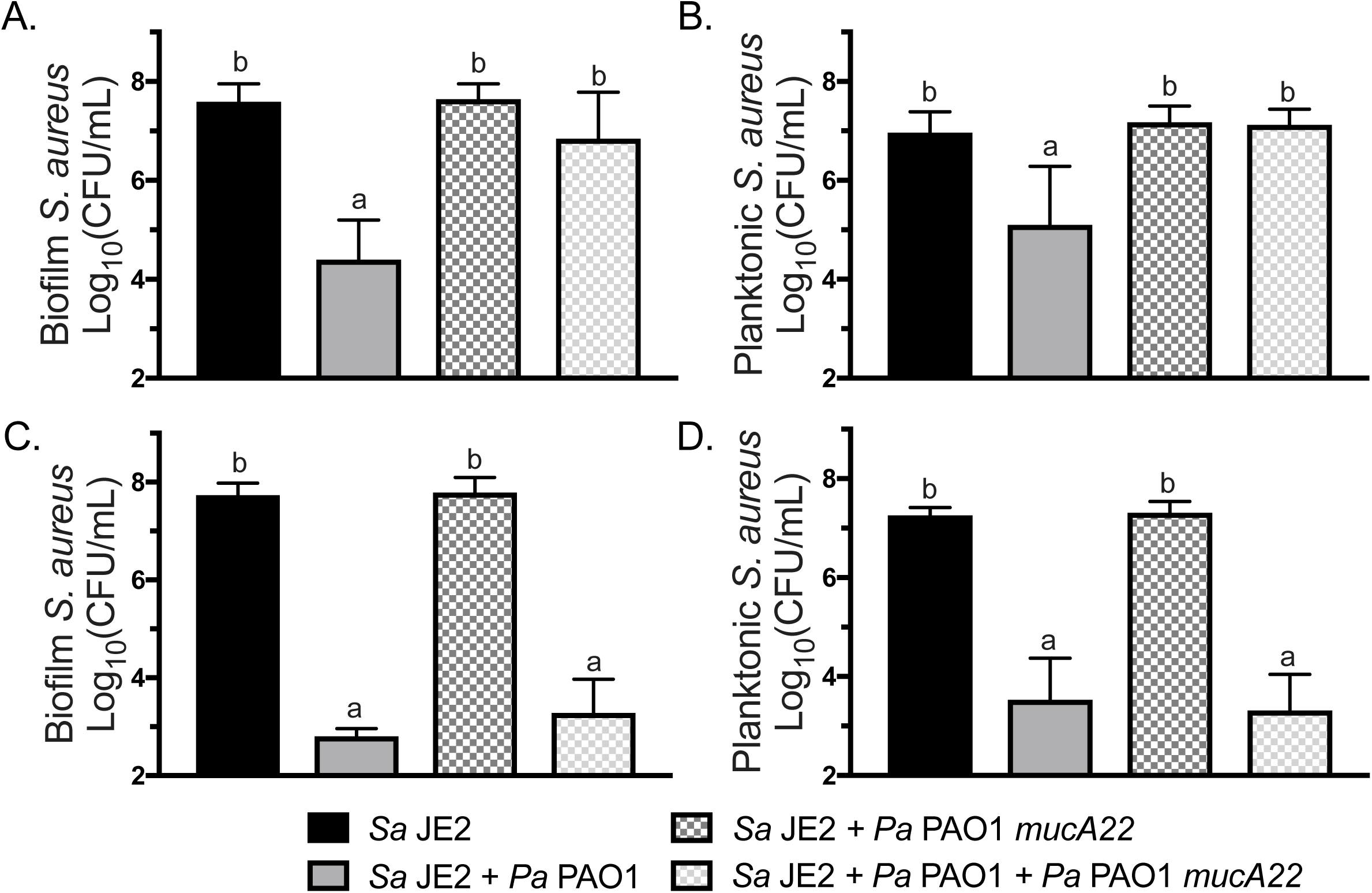
Culture with mucoid *P. aeruginosa* PAO1 *mucA22* delays *S. aureus* JE2 killing by wild-type *P. aeruginosa* PAO1. Biofilm tri-culture on plastic with *S. aureus* JE2, *P. aeruginosa* PAO1, and *P. aeruginosa* PAO1 *mucA22* in MEM L-gln L-arg. CFU/ml were enumerated on MSA and log_10_ transformed. *S. aureus* JE2 viability in the **A)** biofilm and **B)** planktonic fraction after 12 hours and S. aureus JE2 viability in the **C)** biofilm and **D)** planktonic fraction after 16 hours co-culture. Dashed boxes indicate the presence of a mucoid strain in the culture. Significance was determined by one-way ANOVA with Dunnett’s post-test. a, p<0.05 with *S. aureus* JE2 as the reference. b, p<0.05 with *P. aeruginosa* PAO1 + *S. aureus* JE2 as the reference.

### Alginate synthesis is not required for protection by exogenous alginate

We hypothesized that exogenous alginate might protect *S. aureus* by stimulating *P. aeruginosa* to produce alginate and therefore lower virulence towards *S. aureus* via known changes that accompany mucoidy (32). We used a *P. aeruginosa* strain that cannot synthesize alginate, *P. aeruginosa* PAO1 *algD*::FRT, to test whether the ability to self-produce alginate is required for *S. aureus* protection by exogenous alginate. Using the biofilm co-culture model, we measured a significant increase in *S. aureus* survival in the presence of *P. aeruginosa* PAO1 *algD*::FRT with exogenous alginate in both the biofilm and planktonic fractions (Fig. 3 A-B). Growth of *P. aeruginosa* PAO1 and *P. aeruginosa* PAO1 *algD*::FRT was non-significantly and significantly increased, respectively, by exogenous alginate in the biofilm fraction only (Fig. 3 C-D). Therefore, the ability to self-produce alginate is not required to reduce virulence towards *S. aureus* in the presence of exogenous alginate.

**Figure 3.**
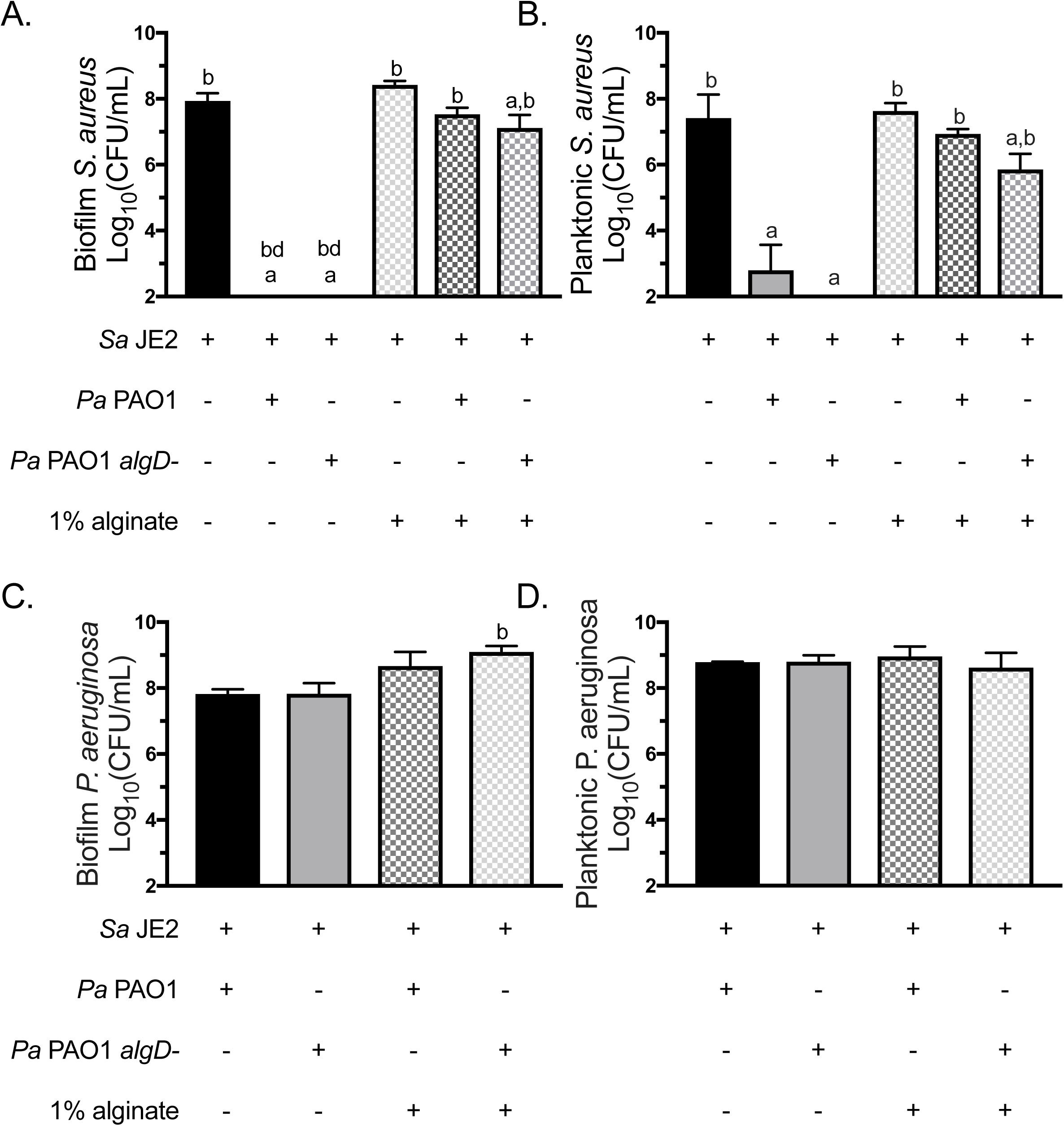
Alginate synthesis is not required for *S. aureus* protection. *S. aureus* JE2 was co-cultured in 1/2X MEM L-gln L-arg on plastic with *P. aeruginosa* PAO1 or *P. aeruginosa* PAO1 *algD*::FRT +/− 1% seaweed-derived alginate. *S. aureus* JE2 viability in the A) biofilm and B) planktonic fractions after 16 hours. *P. aeruginosa* PAO1 viability in the C) biofilm and D) planktonic fractions after 16 hours. Dashed boxes indicate the presence of alginate in the culture. Significance was determined by one-way ANOVA with Dunnett’s post-test. a, p<0.05 with *S. aureus* JE2 as the reference. b, p<0.05 with *P. aeruginosa* PAO1 + *S. aureus* JE2 as the reference.

### Exogenous alginate transcriptionally decreases siderophore production by *P. aeruginosa*

*S. aureus* is protected from mucoid *P. aeruginosa* due to the transcriptional downregulation of genes in the mucoid strain of *P. aeruginosa* required to produce the iron chelator pyoverdine, rhamnolipid surfactant and the redox active molecule HQNO (32). Thus, we investigated whether exogenous alginate might function via a similar mechanism. Pyoverdine production was quantified by measuring relative fluorescence units (RFUs) with excitation at 400 and absorbance at 460 of supernatant from *P. aeruginosa* grown with +/− 1% exogenous alginate*. P. aeruginosa* PAO1 *mucA22* and *P. aeruginosa* PAO1 Δ*pvdA* were included as controls. Pyoverdine decreased significantly when *P. aeruginosa* PAO1 was cultured with exogenous alginate (Fig. 4A). Interestingly, while mucoid *P. aeruginosa* PAO1 *mucA22* produced significantly less pyoverdine than wild-type *P. aeruginosa* PAO1, addition of exogenous alginate further decreased pyoverdine production by the mucoid strain.

**Figure 4.**
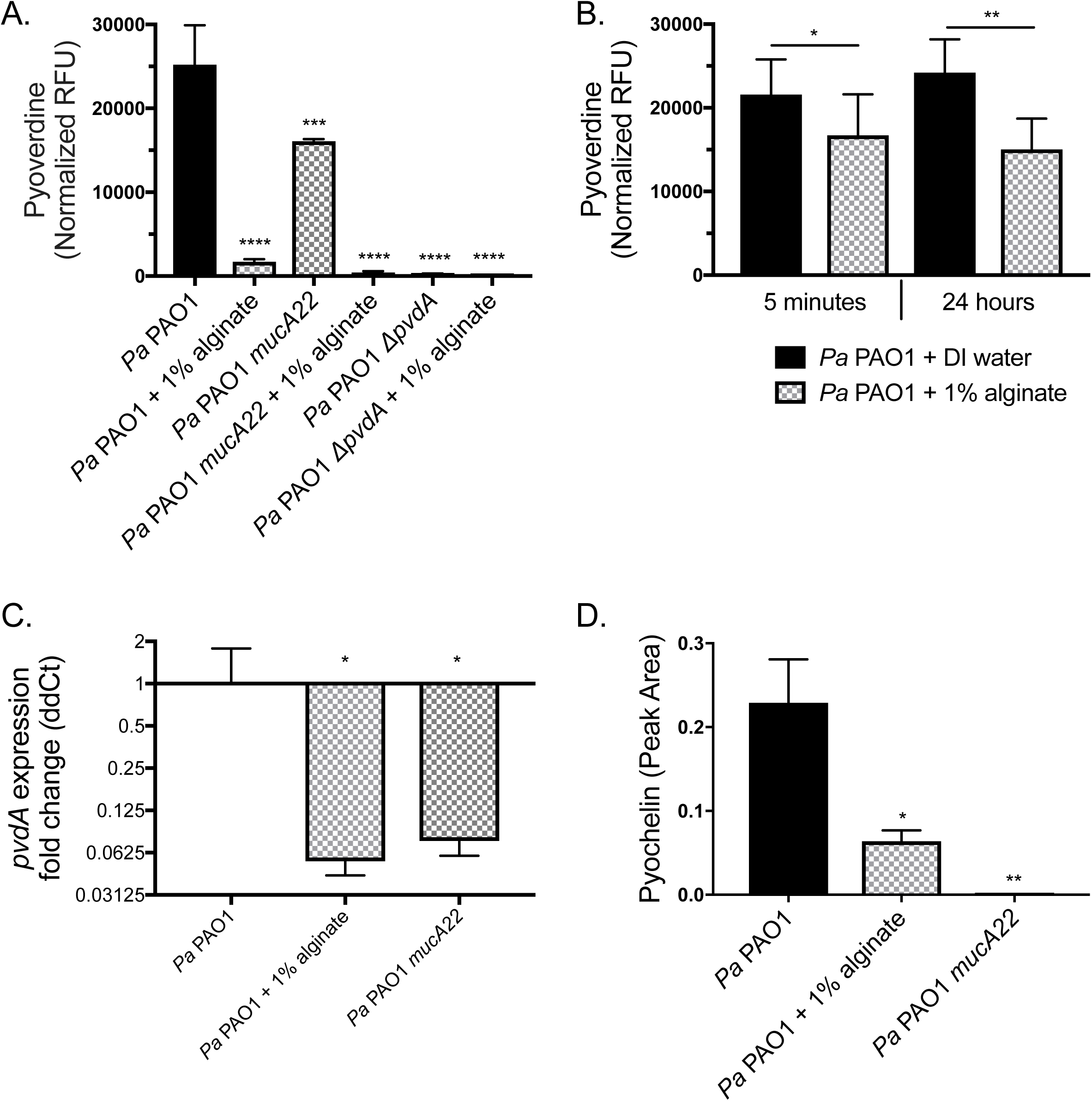
Exogenous alginate decreases *P. aeruginosa* PAO1 siderophore production. **A-B)** *P. aeruginosa* PAO1 was grown on plastic in 1/2X MEM L-gln L-arg +/− 1% alginate for 16 hours at 37°C with 5% CO_2_. Supernatants were collected from the planktonic fraction. **A)** Pyoverdine was quantified by measuring RFU of the supernatants at 400 nm excitation and 460 nm emission and normalizing to CFU/mL of the planktonic fraction. **B)** Supernatants were diluted 1/2X in DI water or 1% alginate and incubated statically at 37°C with 5% CO_2_ for 5 minutes or 24 hours and pyoverdine was quantified by RFU of the supernatants at 400nm excitation and 460nm emission. Significance determined by paired t-test. **C-D)** *P. aeruginosa* PAO1 was grown in 25mL TSB shaking for 8 hours. **C)** *pvdA* expression was quantified by q-RT-PCR and ddCt was calculated relative to *P. aeruginosa* PAO1 *rpoD* expression. **D)** Pyochelin was quantified by LC-MS/MS. Significance determined by one-way ANOVA with Dunnett’s post-test comparison to *P. aeruginosa* PAO1 unless otherwise indicated. For all statistical tests, *p<0.05, **p<0.01, ***p<0.001, ****p<0.0001.

We performed a quenching experiment to test whether exogenous alginate can absorb pyoverdine from the media or mask its signal in our assay. *P. aeruginosa* PAO1 supernatants were diluted 1/2X with DI water or 2% exogenous alginate and pyoverdine was quantified after an additional 5 minute and 24 hour incubation at 37°C. There was a small but significant and repeatable decrease in total pyoverdine measured when supernatants were diluted in alginate versus water (Fig. 4B), indicating that alginate may absorb a small portion of the pyoverdine present or partially mask its signal in our assay.

To test whether pyoverdine production is reduced at the transcriptional level by exogenous alginate, we cultured *P. aeruginosa* PAO1 in the presence and absence of exogenous alginate and measured transcription of *pvdA*, a gene essential for pyoverdine production, by qRT-PCR. The mucoid strain *P. aeruginosa* PAO1 *mucA22* was used as a control, as this strain transcriptionally downregulates *pvdA* (32). The expression of *pvdA* was significantly reduced when *P. aeruginosa* PAO1 was grown with 1% exogenous alginate (Fig. 4C). Consistent with previous data (32), *pvdA* expression was also decreased in the mucoid strain carrying the *mucA22* mutation. When pyoverdine production is low, *P. aeruginosa* will typically increase the production of pyochelin, the other major *P. aeruginosa* siderophore, so we used liquid chromatography tandem mass spectrometry (LC-MS/MS) to quantify pyochelin. *P. aeruginosa* PAO1 also reduced pyochelin production upon exposure to exogenous alginate (Fig. 4D). Taken together, these results indicate that exogenous alginate reduces the production of both *P. aeruginosa* siderophores pyoverdine and pyochelin, likely via transcriptional down regulation, and may also at least partially sequester siderophores that are produced.

### Exogenous alginate post-transcriptionally decreases rhamnolipid production by *P. aeruginosa*

Rhamnolipids are surfactants produced by *P. aeruginosa* that intercalate into the cytoplasmic membrane of *S. aureus* (47). Surfactants disrupt the surface tension of liquids, a characteristic that we used to quantify rhamnolipid production by *P. aeruginosa* using the drop collapse assay (Fig. 5A). We observed a minor, non-significant decrease in total rhamnolipid production by wild-type *P. aeruginosa* PAO1 in the presence of 1% exogenous alginate. The rhamnolipid-deficient strain *P. aeruginosa* PAO1 *rhlA::Gm* was not affected by the presence of exogenous alginate (Fig. 5B).

**Figure 5.**
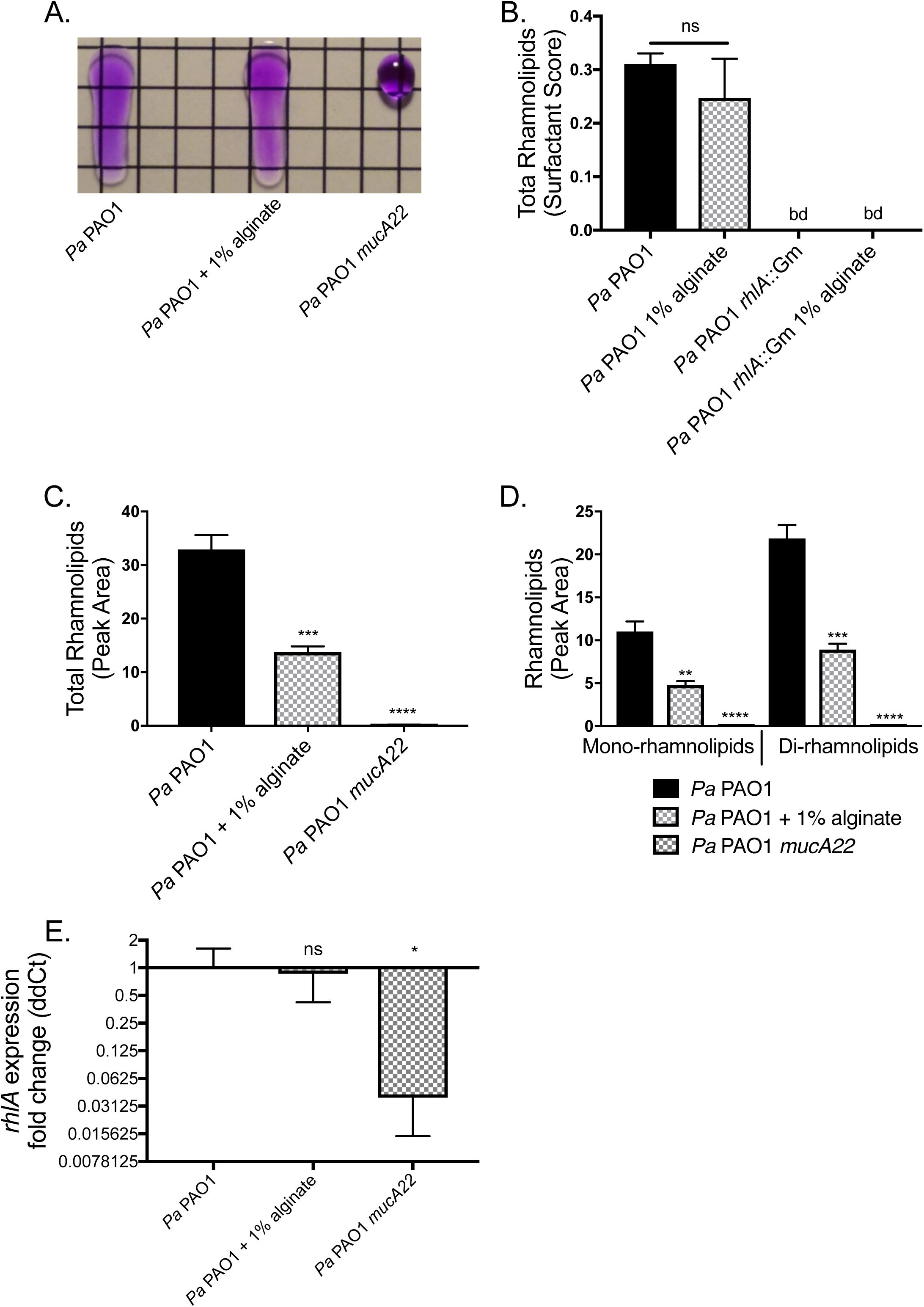
Rhamnolipid production is post-transcriptionally altered by exogenous alginate. *P. aeruginosa* PAO1 was grown in TSB in liquid culture for 8 hours +/− 1% alginate. Supernatants were collected by centrifuging to remove cell debris, and sterile filtering. Cell pellets were snap frozen for expression analyses. **A)** Representative drop collapse assay image. *P. aeruginosa* PAO1 supernatants were prepared from mucoid and non-mucoid *P. aeruginosa* PAO1 and serially diluted 1:2 in PBS. Surfactant activity was assessed by placing a 20uL droplet of each supernatant dilution on plastic, placing the droplet at a 90° angle for 10 seconds and assessing migration (the PBS was supplemented with 0.01% CV to aid visualization). Surfactant activity was quantified as the reciprocal of the highest dilution at which the drop migrates. **B)** Rhamnolipid production by *P. aeruginosa* PAO1 quantified by drop collapse and normalized to CFU/mL to determine surfactant score. Rhamnolipid quantification by LC-MS/MS for **C)** total rhamnolipids and **D)** mono- and di-rhamnolipids. **E)** *rhlA* expression was quantified by q-RT-PCR and ΔΔCt was calculated relative to *P. aeruginosa* PAO1 *rpoD*. Significance determined by one-way ANOVA with Dunnett’s post-test comparison to *P. aeruginosa* PAO1. *p>0.05, **p<0.01, ***p<0.001, ****p<0.0001.

Because the drop collapse assay is an estimate of total rhamnolipids, we also used LC-S/MS to quantify total and specific rhamnolipids in *P. aeruginosa* PAO1 supernatants. Overall rhamnolipid quantity significantly decreased by ∼50% with the addition of exogenous alginate when measured by this more sensitive method (Fig. 5C). The fold-change decrease was similar for both mono- and di-rhamnolipids (Fig. 5D). To determine whether the reduced rhamnolipid production was due to transcriptional regulation, we measured the relative expression of *rhlA* for *P. aeruginosa* PAO1 +/− 1% alginate, and *P. aeruginosa* PAO1 *mucA22*. Transcription of *rhlA* was not significantly different between *P. aeruginosa* PAO1 +/− 1% exogenous alginate (Fig. 5E).

### Exogenous alginate downregulates PQS quorum sensing

The PQS (Pseudomonas quinolone signal; 2-heptyl-3,4-dihydroxyquinoline) quorum sensing system, which impacts a range of *P. aeruginosa* virulence factors (Fig. 6A), is known to be down-regulated by mucoid *P. aeruginosa* strains (32, 48). We therefore measured the impact of exogenous alginate on several products regulated by the PQS quorum sensing system, with *P. aeruginosa mucA22* as a control. Interestingly, both self-produced and exogenous alginate decrease production of several key PQS-regulated products, including HQNO, HHQ (4-hydroxy-2-heptylquinoline), and PQS (Fig. 6B-D) as well as downstream factors such as phenazines PCA (phenazine-1-carboxylic acid) and pyocyanin (Fig. 6E-F).

**Figure 6.**
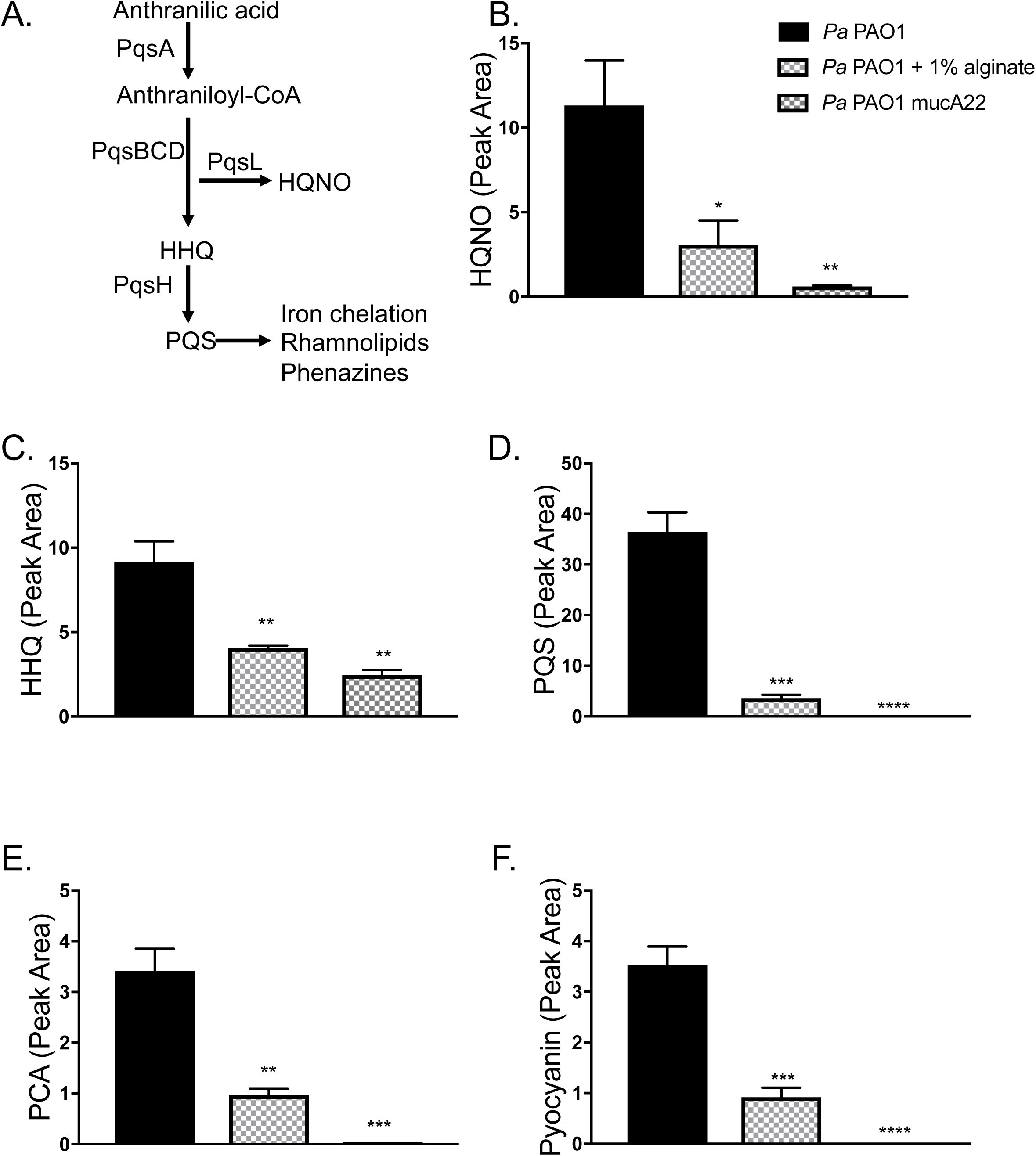
Exogenous alginate downregulates PQS quorum sensing. **A)** The PQS quorum sensing regulon. PQS synthesis is dependent on the *pqsABCD* genes located in the *pqs* operon. Both HHQ and HQNO have direct antimicrobial properties, while PQS is the ligand for PqsR. When PQS level is high this ligand interacts with PqsR to positively regulate many downstream virulence factors. PQS has direct iron-chelating function and promotes expression of siderophore-encoding genes. The downstream effects listed here focus on effects relevant to this study and are not exhaustive. **B-F)** In Panel B is shown is the legend for all graphs in this figure. LC-MS/MS was used to quantify **B)** HQNO **C)** HHQ **D)** PQS **E)** PCA and **F)** pyocyanin produced by *P. aeruginosa*.

### Expression profiles of *P. aeruginosa* in the presence of exogenous alginate

To determine how exogenous alginate impacts *P. aeruginosa* gene expression more generally, we used the Nanostring PAV2 codeset (49) to compare *P. aeruginosa* gene expression in the presence and absence of exogenous alginate and *S. aureus*. Nansotring is a digital multiplexed technology for direct quantification of RNA transcripts. The PAV2 codeset used in this study contains probes for 75 transcripts associated with *P. aeruginosa* genes known or suspected to be expressed in the CF airway (49). *P. aeruginosa* PAO1 and *P. aeruginosa* PA14 were cultured to mid-log phase and then sub-cultured into fresh media +/− 1% alginate. Samples were collected after an additional 45 minutes of growth and expression was analyzed by Nanostring according to the manufacturer’s protocols.

Ten genes were significantly differentially regulated by *P. aeruginosa* PAO1 in the presence of exogenous alginate (Fig. 7A). We previously used the same Nanostring codeset to compare mucoid and nonmucoid *P. aeruginosa* PAO1 gene expression, and measured a significant downregulation of *pvdA, rhlA, pchC, fliC type B*, and *norC*, as well as a significant upregulation of *algI* and *algD.* Of these genes, only *pchC* is significantly differentially regulated in the same direction across both studies, indicating that transcriptional changes due to exogenous alginate are not broadly similar to the effects of self-produced alginate by *P. aeruginosa* PAO1. We observed an overlap in significant transcriptional changes in response to exogenous alginate between *P. aeruginosa* PA14 and *P. aeruginosa* PAO1 for five genes – *cyoA, narG, norC, pchC*, and *pscC* (Fig. 7B). *P. aeruginosa* PA14 also had a large number of genes with a small but significant transcriptional upregulation in the presence of exogenous alginate (Supplemental Tables 1-2).

**Figure 7.**
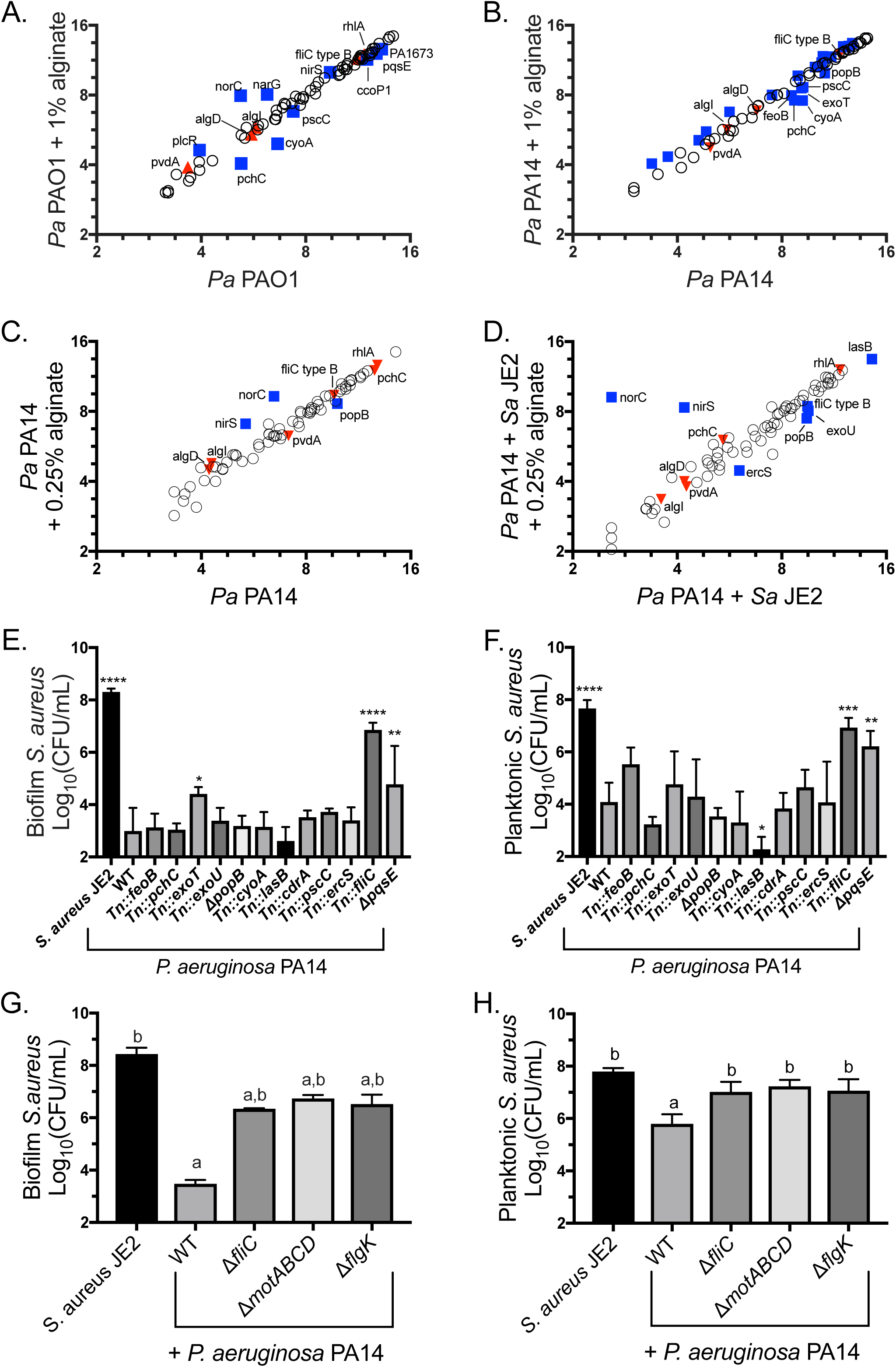
Alginate alters the expression of *P. aeruginosa* genes essential for *S. aureus* killing. **A-D)** Raw Nanostring counts were normalized to positive controls and three housekeeping genes (*rpoD, ppiD, fbp*) and log_2_ transformed. Three biological replicates per condition. Significantly differentially expressed genes were determined by unpaired *t-*test followed by the two-stage linear step-up procedure of Benjamini, Krieger, and Yekutieli (with *q* 1% for false discovery), and are marked by blue squares and labeled with the gene name. Genes significantly differentially expressed by mucoid *P. aeruginosa* in a previous study (32) but not by *P. aeruginosa* in the presence of exogenous alginate are marked by red triangles. **A)** *P. aeruginosa* PAO1 and **B)** *P. aeruginosa* PA14 sub-cultured into TSB or TSB + 1% alginate during mid-log growth phase for 45 minutes. For clarity, only significantly downregulated genes are labeled in **B)**. *P. aeruginosa* PA14 **C)** mono-cultured and **D)** co-cultured with *S. aureus* JE2 in TSB + 0.25% alginate for 8 hours. **E-F)** *S. aureus* JE2 survival in the **E)** biofilm and **F)** planktonic fractions after 16 hour co-culture with *P. aeruginosa* PA14 mutants for genes downregulated in one or more Nanostring experiment. Significance determined by one-way ANOVA with Dunnett’s post-test comparison to *S. aureus* in monoculture. *p<0.05, **p<0.01, ***p<0.001, ****p<0.0001. **G-H)** *S. aureus* JE2 survival in the **G)** biofilm and **H)** planktonic fractions after 16 hour co-culture with *P. aeruginosa* PA14 motility mutants. Significance determined by one-way ANOVA with Dunnett’s post-test. a, p<0.05 with *S. aureus* JE2 as the reference. b, p<0.05 with *P. aeruginosa* PAO1 + *S. aureus* JE2 as the reference.

To understand how longer exposure to exogenous alginate and the presence of *S. aureus* effect *P. aeruginosa* transcription, we cultured *P. aeruginosa* PA14 for 8 hours +/− 0.25% alginate and +/− *S. aureus.* Nanostring counts were normalized and analyzed as previously described (Supplemental Tables 3-4). Only three genes were significantly differentially expressed by *P. aeruginosa* PA14 monocultured in the presence of alginate relative to *P. aeruginosa* PA14 monocultured without alginate *(nirS, norC*, and *popB*; Fig. 7C). When *P. aeruginosa* PA14 was co-cultured with *S. aureus* the same three genes had altered expression, as well as *ercS, exoU, fliC type B, and lasB* (Fig. 7D). Notably, *fliC type B* is also downregulated by mucoid *P. aeruginosa* relative to wild-type *P. aeruginosa* (32). The data was further analyzed by clustering and principal component analysis, with results supporting that *S. aureus* has a larger effect on overall gene expression than exogenous alginate (Fig. S6).

We hypothesized that some of the genes transcriptionally downregulated by *P. aeruginosa* in the presence of exogenous alginate might contribute to *S. aureus* protection in coculture. We focused on downregulated genes because the Nanostring codeset genes are related to virulence and pathogenesis, making it unlikely that upregulation of any of these genes would positively impact *S. aureus* survival. We utilized a *P. aeruginosa* PA14 nonredundant transposon mutant library (50) to test whether any of these factors had a direct impact on *S. aureus* survival. We performed a biofilm coculture experiment for *P. aeruginosa* PA14 mutants of each gene identified as significantly down regulated in any Nanostring experiment and found that disruption of *exoT, fliC*, or *pqsE* significantly enhanced *S. aureus* survival in the biofilm fraction in the presence of *P. aeruginosa* PA14 (Fig. 7E). The *fliC and pqsE* mutants also enhanced *S. aureus* survival in the planktonic fraction (Fig. 7F). Only *fliC* altered *P. aeruginosa* PA14 growth, which was slightly but significantly increased in the biofilm fraction (Fig. S7 A-B).

Because *fliC* mutants do not produce flagella, we suspected functional flagella might be required for *S. aureus* killing. We therefore tested *flgK* and *motABCD* mutants in the biofilm co-culture model and saw that these mutations also prevented *S. aureus* killing by *P. aeruginosa* PA14 (Fig. 7 G-H) without disrupting *P. aeruginosa* PA14 growth (Fig. S7 C-D). Because *flgK* mutants are missing the flagellar cap and *motABCD* mutants produce a nonfunctional flagellum, these findings indicate that flagellar function is required for full virulence of *P. aeruginosa* towards *S. aureus*, and furthermore, suggest that both self-produced and exogenous alginate reduce *P. aeruginosa*-mediated virulence towards *S. aureus*, in part, by downregulating flagellar genes.

### Characterization of mucoid clinical isolates for the ability to kill *S. aureus* in co-culture

Our previous work (23, 32) demonstrated that most mucoid *P. aeruginosa* strains do not kill *S. aureus* when cocultured *in vitro*. Interestingly, other previous work published by our lab indicated that at least one mucoid clinical isolate, *P. aeruginosa* FRD1, can kill *S. aureus* in co-culture (33). To determine whether other mucoid clinical isolates could kill *S. aureus* during *in vitro* co-culture, in part to understand the basis by which these “mucoid killers” could function, we selected a panel of mucoid clinical isolates identified in previous studies (32) to co-culture with *S. aureus*.

We co-cultured *P. aeruginosa* clinical isolates with *S. aureus* JE2 and enumerated the viability of *S. aureus* out to 24 hours of co-culture due to the slow growth of some of the clinical isolates relative to laboratory strains. The ability of mucoid clinical isolates to kill *S. aureus* in coculture varies by strain (Fig. 8A), with some strains such as FRD1 killing robustly, while others such as CFBRPA33 showing no killing of *S. aureus*. Mucoidy does still provide some protection from *S. aureus* even at late time points, as non-mucoid revertants for FRD1 and Cl2224 killed *S. aureus* more efficiently than the mucoid strains at 16 and 24 hours. We quantified reversion rates for *P. aeruginosa* FRD1 and *P. aeruginosa* CFBRPA32 to determine whether killing is due to high rates of reversion to non-mucoid phenotypes, and observed reversion rates <0.1% after 24 hours of culture (Supplemental Table 5).

**Figure 8.**
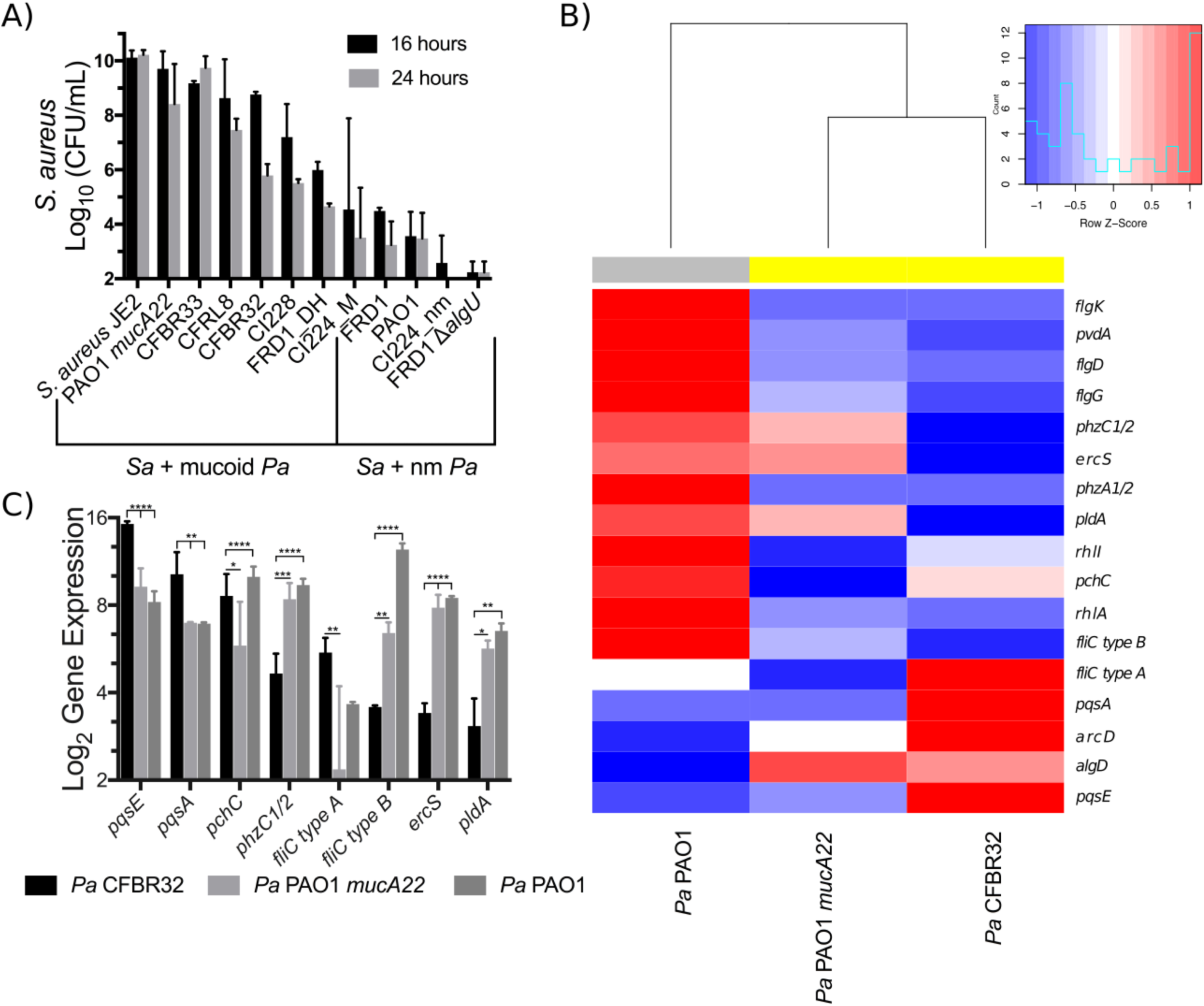
Mucoid clinical isolates have varying effects on *S. aureus* in coculture. **A)** *S. aureus* JE2 growth after 16 and 24 hours of co-culture in flasks shaking at 225 rpm in TSB with *P. aeruginosa* clinical isolates. Nm = nonmucoid. **B-C)** Log_2_ transformation of Nanostring counts normalized to positive controls and three housekeeping genes (*rpoD, ppiD, fbp*) for the indicated transcripts for clinical isolate *P. aeruginosa* CFBRPA32, mucoid laboratory strain *P. aeruginosa* PAO1 mucA22, and wild-type laboratory strain *P. aeruginosa* PAO1 after 24 hours culture in flasks shaking at 225 rpm in TSB. Two biological replicates per strain. Gene expression was analyzed by two-way ANOVA followed by Tukey’s multiple comparisons. **B)** Heatmap and dendrogram of all genes significantly differentially regulated between any two strains. Expression values displayed as within-row Z-scores. Yellow indicates mucoid strains, and gray indicates non-mucoid strains. **C)** All genes significantly differentially regulated between *P. aeruginosa* CFBRPA32 and *P. aeruginosa* PAO1 *mucA22*. *p<0.05, **p<0.01, ***p<0.001, ****p<0.0001.

We used the Nanostring codeset described in the previous section to generate expression data for *P. aeruginosa* PAO1, CFBRPA32 (a “mucoid killer”), and the mucoid laboratory strain *P. aeruginosa* PAO1 *mucA22* after 24 hours of growth. Nanostring counts were normalized as described previously and then analyzed by two-way ANOVA followed by Tukey’s post-test between all three strains (Supplemental Tables 6-7). Of the 75 transcripts measured, seventeen genes were significantly differentially regulated between at least one pair of strains (Fig. 8B).

Clustering by expression of differentially regulated genes revealed that the expression profile of CFBRPA32 is more similar to that of the mucoid laboratory strain *P. aeruginosa* PAO1 *mucA22* than nonmucoid *P. aeruginosa* PAO1 (Fig. 8B). However, eight transcripts were significantly differentially regulated between *P. aeruginosa* CFBRPA32 and *P. aeruginosa* PAO1 *mucA22*: *pqsE, pqsA, pchC, phzC1/2, fliC type A, fliC type B, ercS, and pldA* (Fig. 8C). Increased expression of *pqsE, pqsA* and *pchC* indicates a potential role for both quorum sensing and iron acquisition in the ability of CFBRPA32 to kill *S. aureus*. CFBRPA32 *pqsE* expression is >50-fold higher than *P. aeruginosa* PAO1 *mucA22* and >125-fold higher than *P. aeruginosa* PAO1. Because *pqsE* is essential for *P. aeruginosa* killing of *S. aureus*, high expression may contribute to the ability of *P. aeruginosa* CFBRPA32 to kill *S. aureus* at late time points.

We chose to perform our Nanostring experiment with CFBRPA32 because there is publicly available Nanostring data (49) for several other mucoid clinical isolates we examined (CFRL8, CI2224, CI228, and FRD1). We re-analyzed this data to determine whether we could identify a shared expression pattern between mucoid killers (Fig. S8). However, there was no individual gene other than *algD* that correlated with *S. aureus* survival. Taken together, our data indicate that mucoid *P. aeruginosa* strains with the ability to kill *S. aureus* in co-culture likely do so via strain-specific mechanisms, and furthermore, some of the genes identified in our analysis of “mucoid killers” overlap with functions down-regulated by exogenous alginate.

## Discussion

*P. aeruginosa* and *S. aureus* coexist within the CF lung, where a heterogeneous array of microorganisms, the host immune system, and nutrient availability all contribute to a complex and dynamic environment (8, 13, 31). Here we demonstrate that alginate, which is known to impact both host-pathogen and pathogen-pathogen interactions, also indirectly impacts pathogen-pathogen interactions (51, 52). Protection of *S. aureus* from *P. aeruginosa* by exogenous alginate is consistent across a variety of nutritional and physiological contexts, given that protection occurs in liquid and biofilm modes of growth, and in both rich and minimal media with two strains of *P. aeruginosa* (PAO1 and PA14; Fig. 1; Fig. S1-S4). Furthermore, alginate production by a mucoid strain of *P. aeruginosa* is sufficient to delay *S. aureus* killing by a nonmucoid strain of *P. aeruginosa* when all three strains are cultured together (Fig. 2; Fig. S5), demonstrating the potential for mucoid strains to alter interactions between non-mucoid *P. aeruginosa* and *S. aureus* strains *in vivo*.

While *P. aeruginosa* robustly kills *S. aureus in vitro*, it has been demonstrated that reduction of any single virulence factor will greatly diminish the ability of *P. aeruginosa* to kill *S. aureus*. We demonstrate that production of *P. aeruginosa* siderophores, pyoverdine and pyochelin, is lowered when *P. aeruginosa* is cultured in exogenous alginate and that this decrease is due at least in part to transcriptional downregulation of *pvdA* (Fig. 4). Notably, the decrease in siderophore production in the presence of exogenous alginate may not be solely due to transcriptional changes. The incubation of supernatant from wild-type *P. aeruginosa* with alginate reduces total pyoverdine levels, indicating a possible role for sequestration of siderophores by alginate (Fig. 4A-B). Exogenous alginate, like self-produced alginate, causes transcriptional downregulation of *pvdA* expression, indicating that self-produced alginate may also lead to decreased expression of this gene via an outside-in signaling mechanism. There is precedence for exogenous exopolysaccharides regulating transcription, as the exopolysaccharide Psl is known to act as an extracellular signal to increase biofilm formation by stimulating c-di-GMP synthesis, although the mechanism is still unknown (53). We also demonstrate a modest decrease in total rhamnolipids across multiple culture methods which is not dependent on transcriptional regulation (Fig. 5). However, it is unclear whether this modest rhamnolipid decrease is enough to affect *S. aureus* survival. Exogenous alginate also interferes with *P. aeruginosa* quorum sensing, as demonstrated by the decrease in levels of HQNO, AQs, and phenazines (Fig. 6). The combined interference with HQNO, siderophore, and phenazine production all likely contributes to *S. aureus* survival in the presence of exogenous alginate. Finally, we demonstrate that a functional flagellum is required for *S. aureus* killing. While flagella are considered virulence factors in the host-pathogen context, their role in *P. aeruginosa-S. aureus* interactions has not previously been demonstrated. Furthermore, this demonstrates that exogenous alginate has the capacity, at least in part, to affect *P. aeruginosa* strains in a manner analogous to self-produced alginate, that is by reducing production of anti-*Staphylococcal* factors. An interesting question that remains is whether the response of *P. aeruginosa* to exogenous alginate is specific; it is possible that the mechanism by which *P. aeruginosa* senses and responds to alginate is due, for example, to increased viscosity. Alternatively, alginate could be enforcing spatial structure or influencing *S. aureus* gene expression.

Interestingly, the presence of alginate can be overcome in some mucoid clinical isolates through unique re-wiring of gene expression (Fig. 8), with the apparent increased expression of genes coding for one or more key anti-*Staphylococcal* factors. Thus, endogenous and exogenous alginate seem to impact an overlapping set of functions that modulate the interaction between *P. aeruginosa* and *S. aureus*. Our observations are not entirely surprising in the context of the evolving nature of both *P. aeruginosa* and *S. aureus* within the CF lung. Finally, our study highlights the importance of the physical environment within the CF lung on the outcomes of polymicrobial interactions, and the need for further studies examining how different specific aspects of the CF lung environment can impact both polymicrobial interactions and overall patient outcomes.

## Materials and Methods

Detailed protocols and strains used in this study are available in the supplemental text.

### Media

All experiments were performed in tryptic soy broth (TSB) or MEM. MEM was supplemented with 2 mM L-glutamine (L-gln) +/− 0.4% L-arginine (L-arg). For experiments containing alginate, MEM was diluted ½X with sterile DI water or alginate. Seaweed alginate was prepared by dissolving alginic acid sodium salt (Sigma) in DI water and autoclaving to sterilize.

### Cocultures

Overnight cultures were diluted to a final concentration of 0.05 OD. For experiments in MEM, strains were centrifuged and resuspended in MEM L-Gln prior to dilution. Strains in flasks were grown shaking at 225rpm at 37°C, and tubes were placed on a culture wheel at 37°C. 100uL total volume was inoculated in a plastic 96-well plate for biofilm cultures. At indicated time points, culture samples were serially diluted in PBS, and plated onto Pseudomonas Isolation Again (PIA) or Mannitol Salt Agar (MSA) to quantify viable *P. aeruginosa* and *S. aureus*, respectively.

### Alginate preparation and quantification

*P. aeruginosa* PAO1 *mucA22* was cultured in 25mL TSB overnight shaking at 225 RPM and 37°C and alginate was isolated as previously described (32), with minor modifications detailed in the supplemental methods. The concentration of alginate in solution was determined by the carbazole method described by Knutson and Jeanes (54).

### Siderophore quantification

Relative fluorescence of supernatants were quantified at 400nm excitation and 460nm emission. Measurements were normalized to CFU/mL.

### Drop collapse assay

Supernatants were serially diluted ½X in PBS and were scored as the reciprocal of the highest dilution at which the drop collapsed. Scores were normalized to CFU/mL.

### q-RT-PCR

Cell cultures were pelleted (centrifuge 2 min, 14,000 rpm at 4°C) and immediately frozen in ethanol cooled with dry ice. RNA was isolated with TRIzol (Zymo Research) as described by the manufacturer. DNA was removed by three sequential treatments with Turbo-DNAse treatment (Invitrogen), cDNA was prepared (RevertAid First Strand cDNA synthesis kit), and qRT-PCR was performed with SsoFast EvaGreen Supermix (BioRad).

### LC-MS/MS

Detailed descriptions of LC-MS/MS supernatant extraction, analysis, feature finding, and molecular annotation are available in the supplemental text.

### Nanostring

Total RNA was prepared as described for q-RT-PCR. The PAV2 codeset (49) was incubated with total RNA as described by the manufacturer’s protocol.

## Acknowledgements

This work was supported by NIH Grant R37 AI83256-06 to G.A.O and T32AI007363 to C.E.P. Additional support was provided by the CF-Research Development Program (STANTO07R0), DartCF and the BBC (P30-DK117469). The Phelan lab is supported by the ALSAM Foundation (L.S. Skaggs Professorship and Therapeutic Innovation Award) and the NIH (R35 GM128690-01). We thank D.A. Hogan for providing access to clinical isolates, AE Baker for PA14 Δ*fliC*, and Tom Hampton for providing advice on expression data analysis. Clinical CF isolates CFBRPA32 and CFBRPA33 were provided by the CF Biospecimen Registry at the Children’s Healthcare of Atlanta and Emory University CF Discovery Core.

## Figure legends

**Figure S1. Exogenous alginate protects *S. aureus* JE2 from *P. aeruginosa* PAO1 in co-culture. A)** *S. aureus* JE2 and **B)** *P. aeruginosa* PAO1 growth after 8 hours of liquid co-culture in TSB +/− 1% seaweed-derived or *P. aeruginosa* PAO1-derived alginate. CFU/ml were enumerated and log_10_ transformed. Dashed boxes indicate the presence of alginate in the culture. Significance was determined by one-way ANOVA with Dunnett’s post-test. a, p<0.0001 with *S. aureus* JE2 as the reference. b, p<0.0001 for panel **A** and p<0.05 for panel **B** with *P. aeruginosa* PAO1 + *S. aureus* JE2 as the reference.

**Figure S2. Exogenous alginate protects *S. aureus* JE2 from *P. aeruginosa* in co-culture.** Liquid co-cultures of *S. aureus* JE2 and *P. aeruginosa* in TSB +/− seaweed-derived alginate added at 0.25% or 2%. CFU/ml were log_10_ transformed. **A)** *S. aureus* JE2 and **B)** *P. aeruginosa* PAO1 growth curves over 8 hours in monoculture or the indicated cocultures. CFU/ml were enumerated at indicated time points. **C)** *S. aureus* JE2 and **D)** *P. aeruginosa* PAO1 growth after 8 hours. **E)** *S. aureus* JE2 and **F)** *P. aeruginosa* PA14 growth curves over 8 hours. CFU/ml were enumerated at indicated time points. **G)** *S. aureus* JE2 and **H)** *P. aeruginosa* PA14 growth after 8 hours. Dotted lines/dashed boxes indicate the presence of alginate in the culture. Significance was determined by one-way ANOVA with Dunnett’s post-test. a, p<0.05 with *S. aureus* JE2 as the reference. b, p<0.001 with *P. aeruginosa* + *S. aureus* JE2 as the reference.

**Figure S3. Exogenous alginate protects *S. aureus* JE2 from *P. aeruginosa* in co-culture. A-D)** Liquid co-cultures of *S. aureus* JE2 and *P. aeruginosa* PAO1 in TSB +/− seaweed-derived alginate added at 0.25% or 2%. *P. aeruginosa* initial density was 10X higher than *S. aureus* density based on OD_600_. CFU/ml were log_10_ transformed. **A)** *S. aureus* JE2 and **B)** *P. aeruginosa* PAO1 growth curves over 8 hours. CFU/ml were enumerated at indicated time points. The dashed line at 2 indicates the limit of detection. **C)** *S. aureus* JE2 and **D)** *P. aeruginosa* PAO1 growth after 8 hours. **E-F)** Biofilm co-cultures of *P. aeruginosa* PAO1 and *S. aureus* JE2 for 16 hrs in MEM L-gln L-arg +/− 1% seaweed-derived alginate. CFU/ml were enumerated and log_10_ transformed. *P. aeruginosa* PAO1 **E)** biofilm and **F)** planktonic growth. **G-J)** Biofilm co-cultures of *P. aeruginosa* PA14 and *S. aureus* JE2 for 16 hours in MEM L-gln L-arg +/− 1% seaweed-derived alginate. CFU/ml were enumerated and log_10_ transformed. *S. aureus* JE2 **G)** biofilm and **H)** planktonic and *P. aeruginosa* PA14 **I)** biofilm and **J)** planktonic growth. Dotted lines/dashed boxes indicate the presence of alginate in the culture. Significance was determined by one-way ANOVA with Dunnett’s post-test. a, p<0.05 with *S. aureus* JE2 as the reference. b, p<0.05 with *P. aeruginosa* + *S. aureus* JE2 as the reference.

**Figure S4. Exogenous alginate protects *S. aureus* JE2 from *P. aeruginosa* in minimal medium liquid co-culture. A)** *S. aureus* JE2 and **B)** *P. aeruginosa* PAO1 and *P. aeruginosa* PA14 growth after 20 hours co-culture in 1/2X MEM L-gln L-arg +/− 1% alginate. CFU/ml were enumerated and log_10_ transformed. Significance determined by unpaired t-test. *p<.05, **p<.01, ***p<.001.

**Figure S5. Culture with mucoid *P. aeruginosa* PAO1 *mucA22* delays *S. aureus* JE2 killing by wild-type *P. aeruginosa* PAO1.** Biofilm tri-culture on plastic with *S. aureus* JE2, *P. aeruginosa* PAO1, and *P. aeruginosa* PAO1 *mucA22* in MEM L-gln L-arg. CFU/ml were enumerated on MSA and log_10_ transformed. Growth curve of *S. aureus* JE2 viability in the **A)** biofilm and **B)** planktonic fraction. Dashed lines indicate the presence of a mucoid strain in the culture. *P. aeruginosa* PAO1 growth in the **C)** biofilm and **D)** planktonic fraction after 12 hours and *P. aeruginosa* PAO1 growth in the **E)** biofilm and **F)** planktonic fraction after 16 hours co-culture. Dashed boxes indicate the presence of a mucoid strain in the culture. Significance was determined by one-way ANOVA with Dunnett’s post-test. a, p<0.05 with *S. aureus* JE2 as the reference. b, p<0.05 with *P. aeruginosa* PAO1 + *S. aureus* JE2 as the reference.

**Figure S6. *P. aeruginosa* PA14 gene expression changes in response to *S. aureus* JE2 and alginate.** Transcriptional profiling of *P. aeruginosa* by Nanostring revealed that the presence of *S. aureus* has a stronger impact on overall *P. aeruginosa* gene expression than exogenous alginate. *P. aeruginosa* PA14 was cultured for 8 hours in TSB +/− 0.25% alginate and +/− *S. aureus* JE2. Nanostring counts were normalized to positive controls and three housekeeping genes (*rpoD, ppiD, fbp*) and log_2_ transformed. *P. aeruginosa* PA14 gene expression was driven more by exposure to *S. aureus* JE2 than exposure to exogenous alginate. **A)** Clustering by Euclidean distance of the average of three biological replicates. Image was generated in R with the Heatmap.2 function. **B)** Principal component analysis of all biological replicates. Samples separate more strongly on the presence of *S. aureus* (PCA 1) than on the presence of exogenous alginate (PCA 2).

**Figure S7. *P. aeruginosa* PA14 mutants do not have growth defects.** Growth of *P. aeruginosa* PA14 mutants after 16 hours co-culture on plastic with *S. aureus* JE2 in MEM L-gln L-arg. **A)** Biofilm and **B)** planktonic fractions for initial mutant screen. **C)** Biofilm and **D)** planktonic fractions for flagellar mutant screen. Significance determined by one-way ANOVA with Dunnett’s post-test comparison to wild-type *P. aeruginosa* PA14. *p<0.05.

**Figure S8. Gene expression of *P. aeruginosa* clinical isolates and nonmucoid revertants.** Heatmap and dendrogram of a subset of genes in the Nanostring codeset from *P. aeruginosa* clinical isolates cultured in lysogeny broth (LB) for 8-11 hours (Gifford AH, Willger SD, Dolben EL, Moulton LA, Dorman DB, Bean H, Hill JE, Hampton TH, Ashare A, Hogan DA. 2016. Infect Immun 84:2995–3006.). Log_2_ transformation of Nanostring counts were normalized to positive controls from a single biological replicate. Expression values are displayed as within-row Z-scores. Yellow indicates mucoid strains, and gray indicates non-mucoid strains. Image generated in R with Heatmap.2 function. This dataset contains expression data for the isogenic nonmucoid revertants of *P. aeruginosa* FRD1 and *P. aeruginosa* CI2224. Here we display a direct comparison of gene expression for these six clinical isolates and two laboratory *P. aeruginosa* strains of a subset of genes in the Nanostring codeset. Interestingly, isolates cluster primarily on mucoidy, and the mucoid isolates which were better able to kill *S. aureus* (*P. aeruginosa* FRD1 and *P. aeruginosa* CI228) clustered together within those groups. Mucoid *P. aeruginosa* FRD1 decreases some genes as expected given its mucoid phenotype, including *flgG, flgD, flgK*, and *fliC.* However, *rhlA, phz*, and *pvdA* expression remain high; all of these genes are known to contribute to the ability of *P. aeruginosa* to kill *S. aureus* in co-culture (Limoli DH, Hoffman LR. 2019. Thorax thoraxjnl-2018-212616.). *P. aeruginosa* CI228 has a similar gene expression pattern to *P. aeruginosa* FRD1, but *pvdA* expression is even higher in this strain. *P. aeruginosa* CI2224, which is similar to *P. aeruginosa* CFBRPA32 in the ability to kill *S. aureus* (that is, both strains show limited ability to kill *S. aureus*) downregulates a large number of genes coding for known virulence factors, including those genes required for the production of rhamnolipids (*rhlA, rhlI*), phenazines (*phzA, phzC*), quorum sensing factors (*pqsH, pqsA*), and flagella (*flgD, flgG, fliC*). It is not immediately clear based on these data why this isolate is able to kill *S. aureus* at the late time points. *P. aeruginosa* CFRL8, which is the poorest of these four strains at killing *S. aureus*, highly expresses *pchC* and *fliC*, but has lower *pqsA* expression relative to the other strains. To determine whether we could broadly correlate the expression of any particular genes with changes in virulence towards *S. aureus*, we plotted the expression of each gene against *S. aureus* survival in co-culture at both 16 and 24 hours for each *P. aeruginosa* strain where we have Nanostring data and performed a linear regression (data not shown). We observed by this method that no individual gene’s expression is significantly correlated with *S. aureus* survival. Taken together, these data suggest that across clinical mucoid strains, the relative production of a variety of transcripts such as those encoding pyoverdine, rhamnolipids, flagella, and phenazines determines whether isolates of *P. aeruginosa* effectively outcompete *S. aureus*, but the specific mechanisms vary on a strain-by-strain basis.

